# Polarized endosome dynamics engage cytosolic Par-3 and dynein during asymmetric division

**DOI:** 10.1101/2020.05.15.098012

**Authors:** Xiang Zhao, Kai Tong, Xingye Chen, Bin Yang, Qi Li, Zhipeng Dai, Xiaoyu Shi, Ian B. Seiple, Bo Huang, Su Guo

## Abstract

Asymmetric cell division (ACD), which produces two daughters with different fates, is fundamental for generating cellular diversity. In the developing embryos of both invertebrates and vertebrates, asymmetrically dividing progenitors generate daughter cells with differential activity of Notch signaling^1–7^, a key regulator of cell fate decisions^8,9^. The cell polarity regulator Par-3 is critical for establishing this Notch asymmetry^1,4,6^, but the underlying mechanisms are not understood. Here, employing *in vivo* time-lapse imaging in the developing zebrafish forebrain during the mitotic cycle of radial glia, the principal vertebrate neural stem cells^10,11^, we show that during ACD, endosomes containing the Notch ligand Delta D (Dld) undergo convergent movement toward the cleavage plane, followed by preferential segregation into the posterior (and subsequently basal) Notch^hi^ daughter. This asymmetric segregation requires the activity of Par-3 and the dynein motor complex. Employing label-retention expansion microscopy, we further detect Par-3 in the cytosol in association with the dynein light intermediate chain 1 (DLIC1) on Dld endosomes, suggesting a direct involvement of Par-3 in dynein-mediated polarized transport of Notch signaling endosomes. Our data reveal an unanticipated mechanism by which Par-3 regulates cell fate decision by directly polarizing Notch signaling components during ACD.

## Main

Progenitor cells need to properly balance self-renewal and differentiation. Asymmetric cell division (ACD) is an important means to impart these distinct potentials to daughter cells. In metazoans, Partitioning-defective (Par) protein complexes, originally discovered in *C. elegans*^12^, are evolutionarily conserved regulators of cell polarity and ACD^13–17^. Among them, Par-3 (also called PARD3, Bazooka) has been studied in the context of neural progenitor self-renewal and differentiation from *Drosophila* to mammals^18–24^. In vertebrate radial glia progenitors (RGPs), Par-3 is localized to the apical cell membrane and is required to establish in daughter cells the asymmetric activity of Notch signaling (Notch^hi^ vs. Notch^lo^)^4,6,25,26^. Despite these advances, it is not known how Par-3, thought to regulate polarity exclusively through its oligomeric scaffolding properties at the apical or anterior cell membrane^27^, leads to differential Notch activity in the nuclei of daughter cells.

### *In vivo* time-lapse imaging reveals polarized dynamics of Notch ligand-containing endosomes during RGP ACD

As shown previously, in the developing zebrafish forebrain, a majority of RGPs undergo ACD to generate an apical differentiating daughter with low Notch activity and a basal self-renewing daughter with high Notch activity^4^. To understand how such Notch signaling asymmetry arises, we visualized internalized Notch ligand DeltaD (Dld) employing an antibody uptake assay^28^ and *in vivo* time-lapse imaging. Punctate labeling was observed in RGPs (Fig. 1a). Such labeling was lost in the *mind bomb* (*mib*) mutant (Extended Data Fig. 1a), which disrupts a conserved ubiquitin E3 ligase essential for Notch ligand endocytosis^29^. These data indicate that the punctate structures are internalized Dld in endosomes (in short, Dld endosomes). Additionally, we evaluated whether this labeling method impacted embryonic development or RGP cell division modes. No significant effects were observed on these processes (Extended Data Fig. 1b-d).

**Figure 1.**
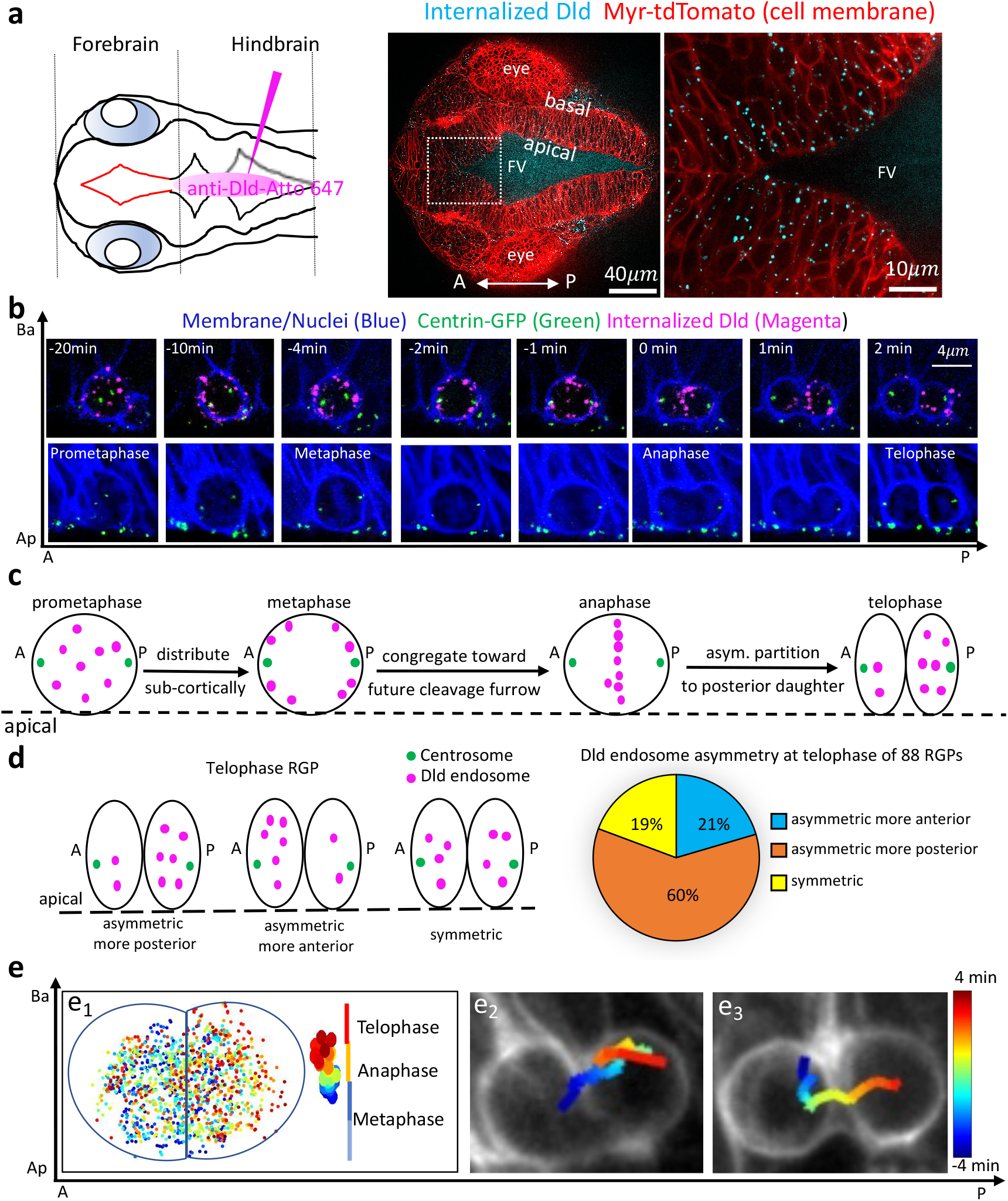
*In vivo* time-lapse imaging reveals polarized dynamics of Notch ligand-containing endosomes during radial glia asymmetric cell division. **a**, Schematic diagram of a 28 hours post fertilization (hpf) embryo (dorsal view) indicating the site of anti-Delta D-Atto 647N (Dld) antibody injection (left). (right) Internalized Dld (blue) imaged 2h after antibody injection shows punctate appearance in RGPs (membrane labeled red). FV, Forebrain ventricle. **b**, Time-lapse imaging of mitotic RGP showing the dynamics of labeled Dld endosomes. Membrane is marked with Myr-TdTomato and nucleus is marked with H2B-mRFP (both pseudo-colored blue), while centrosomes are marked with GFP-Centrin. Time 0 denotes the onset of anaphase (determined by nuclear position and first appearance of the cleavage furrow). **c**, Schematic diagram showing typical phases of internalized Dld dynamic patterns (from 30 RGPs imaged). **d**, Schematic showing the distribution patterns of Dld endosomes at the telophase of mitotic RGPs (See Methods for the quantification of asymmetric index). **e**, Plots showing automated Dld endosome tracking. **e** 1, From a composite of 19 RGPs, each with supernumerary labeled endosomes and complete time-lapse data from prometaphase to telophase. Each dot represents a tracked endosome at a given time, and color codes for cell cycle stages. **e2** and **e3**, two mitotic RGPs each with a single labeled Dld endosome, tracked from −4 min to 4 min (Time is color-coded, and 0 min indicates the onset of anaphase). A-P indicates anterior-posterior axis, and Ap-Ba indicates apical-basal axis.

To observe *in vivo* Dld endosome dynamics during RGP divisions, we performed time-lapse imaging using 24-30 hours post fertilization (hpf) *Tg[ef1a-MyrTdTomato]* embryos (marking cell membranes); the centrosomes were labeled by microinjection of the *GFP-centrin* mRNA. The cell cycle stage of dividing RGPs was determined using *Tg[ef1a-MyrTdTomato; ef1a-H2BmRFP]* embryos, which marked both the cell membrane and nucleus, enabling a correlation between cell shape and DNA patterns (Fig. 1b, time 0 represents anaphase when incoming cleavage furrows become first visible). During imaging, both the apical-basal and the anterior-posterior axes of each RGP were tracked. By analyzing these dynamic videos (See Supplementary Video 1), we made several intriguing observations (Fig. 1b-c, and Extended Data Fig. 2). Dld endosomes, apparently entering RGPs from surrounding cell membranes, were distributed throughout the cytosol during prophase to prometaphase. During metaphase, most Dld endosomes were distributed sub-cortically. By anaphase, however, most Dld endosomes congregated toward the future cleavage plane, and subsequently were unequally partitioned into the posterior daughter after division. An analysis of 88 RGPs at telophase showed that a majority (60%) of them asymmetrically partitioned Dld endosomes into the posterior daughter. Some RGPs with symmetric or anteriorly enriched Dld endosomes were also observed (Fig. 1d; see Methods for the quantification of asymmetry index). Despite such differences, these observed RGPs were distributed around the forebrain ventricle in an intermingled manner (Extended Data Fig. 3).

Automated tracking through the entire mitotic cycle of 30 RGPs allowed us to visualize the progressive enrichment of endosomes toward the posterior daughter at telophase (Fig. 1e_1_, and Supplementary Video 2). Such enrichment could be due to directional endosome movement toward the posterior, their selective degradation at the anterior, or both. The presence of supernumerary labeled Dld endosomes made it challenging to unambiguously discern individual endosome’s trajectories. Intriguingly, due to the mosaic nature of Dld labeling method, some RGPs contained only a single labeled Dld endosome. This enabled us to clearly track the movement of individual endosomes. We observed that the Dld endosome first moved toward the cleavage plane, followed by directed maneuver toward the posterior side (Fig. 1e2-e3, Supplementary Videos 3-6). Together, these data uncover polarized dynamics of Dld endosomes during RGP division and show that in most cases, they are asymmetrically segregated into the posterior daughter after division.

### Notch ligand-containing endosomes preferentially segregate to the posterior-then-basal Notch^hi^ daughter

We next asked what is the possible fate of the daughter that receives a higher amount of Dld endosomes: will it be Notch^hi^ with more self-renewing potential or Notch^lo^ with more differentiation potential? Based on the conventional thinking, that is, a cell with higher amount of Notch ligand activates more Notch receptor in its neighbor(s), one would expect that the daughter with more Dld will be Notch^lo^. In contrast, a previous study employing the *Drosophila* sensory organ precursor (SOP) system has uncovered the co-presence of Delta and Notch in the same endosomes^1^. If this were the case, one would predict that the daughter with higher amount of Dld would also have higher amount of Notch activity (i.e. Notch^hi^)(Fig. 2a).

**Figure 2.**
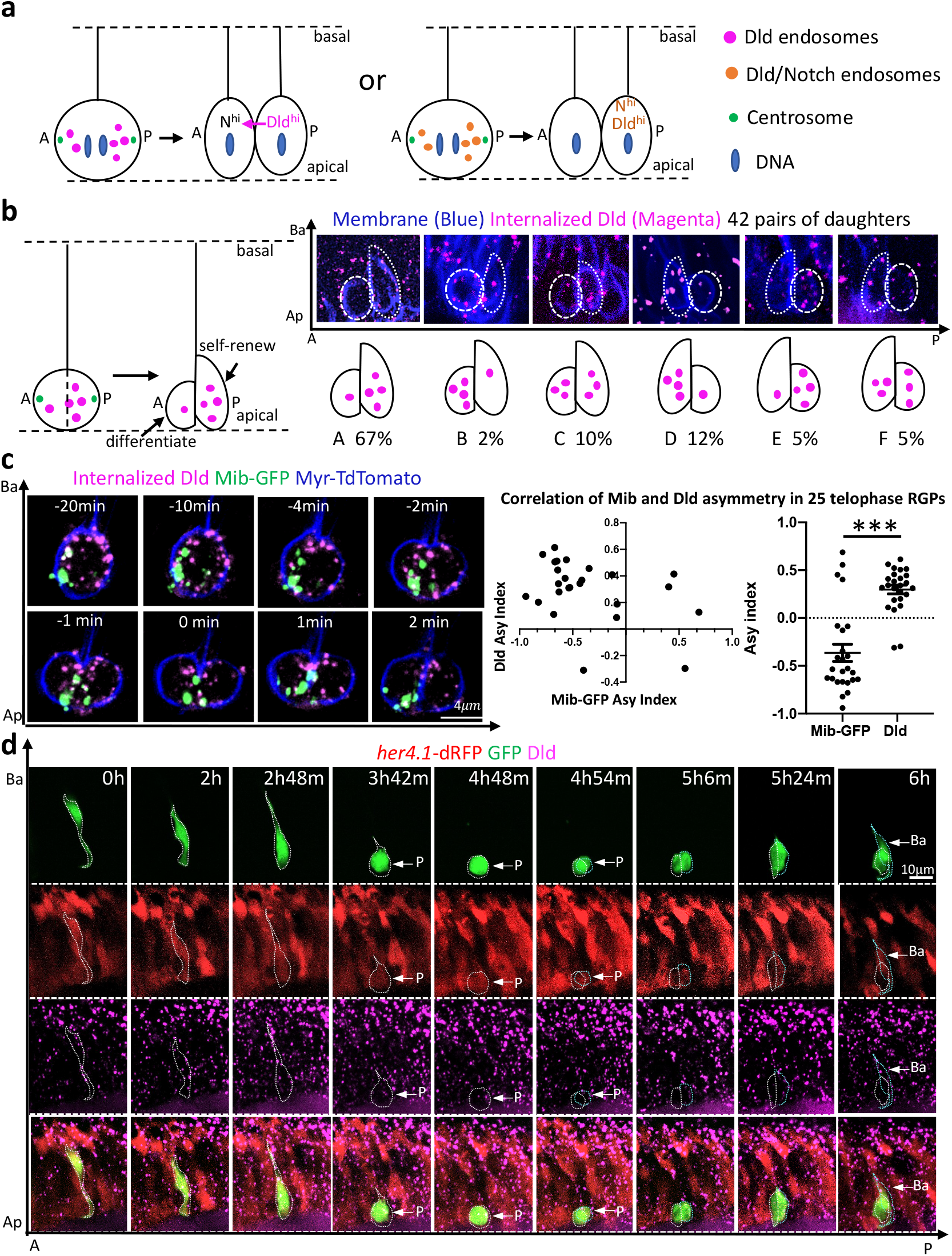
Dld endosomes are preferentially segregated into the Notch^hi^ daughter of asymmetrically dividing RGPs. **a**, Schematic showing two models of potential fate outcomes of daughter cells, Notch^hi^ vs. Notchlo, following asymmetric segregation of Dld endosomes. **b**, Topology and statistics of relative daughter cell position along A-P (anterior posterior) and Ap-Ba (apical basal) axes. Majority of dividing RGPs preferentially segregate Dld endosomes into the posterior (and subsequently basal) daughter (A, 67%, n=42). **c**, Time-lapse sequence of images (left) and statistical plots (right) show the dynamics and preferential segregation of Dld endosomes and Mib into different daughter cells of dividing RGPs (n=25). The left graph plots individual RGP’s asymmetry indices for Mib-GFP (X-axis) and internalized Dld (Y-axis), with the correlation coefficient r = −0.39 (anti-correlation). The right graph shows the distribution of asymmetry indices, with negative values indicating enrichment at the anterior. *P*< 0.0001, t = 6.549, df = 48, unpaired two tailed t-test. The 25 RGPs were derived from 8 embryos of 5 repeat experiments. **d**, Time-lapse sequence of images showing a clonally labeled RGP (green), which divides to preferentially segregate internalized Dld into the daughter cell with higher *her4.1-dRFP* (Ntoch^hi^). Similar patterns of segregation were observed in 3 out of 5 RGPs imaged. The other two RGPs showed symmetric *her4.1-dRFP* distribution in daughter cells after division.

Due to the lack of an anti-Notch antibody that works in zebrafish, we took three different approaches to address possible fate outcomes of daughter cells following RGP division. First, most RGPs that we imaged in the developing zebrafish neurogenic forebrain undergo horizontal divisions along the anteroposterior (A-P) axis. Shortly after division, daughter cells begin interkinetic nuclear migration (INM), and adopt differential position along the apicobasal (Ap-Ba) axis. The basal daughter is previously shown to be Notch^hi^ with higher self-renewing potential^4^ (Fig. 2b). We therefore determined the relationship between the A-P daughter position immediately following RGP mitosis and the Ap-Ba daughter position shortly thereafter. A majority of the nascent posterior daughter (67%, n=42) were found to initiate an earlier basal migration (Fig. 2b). This suggests that Dld endosomes are preferentially segregated to the posterior daughter, which later becomes the basal Notch^hi^ daughter.

The second approach involved analysis of the E3 ubiquitin ligase Mib, which is asymmetrically segregated to the apical differentiating daughter following RGP ACD in zebrafish^4^. Similar observations are also reported in the chick neural progenitors^30^. By simultaneously tracking Dld endosomes and Mib distribution, we found that they were segregated into different daughter cells (Fig. 2c, Extended Data Fig. 4, Supplementary Video 7). This observation indicates that Dld endosomes preferentially segregate away from the Mib-high apical differentiating daughter.

Finally, we performed *in vivo* time-lapse imaging of internalized Dld in *Tg[her4.1-dRFP]*^31^ embryos, which express the Notch activity reporter (*her4.1-dRFP*) in RGPs. The results showed that the posterior daughter with more Dld endosomes later assumed a more basal position with higher *dRFP* fluorescence (i.e. higher Notch activity)(Figure 2d and Extended Data Fig. 5, Supplementary Video 8). Taken together, our findings indicate that the daughter cell receiving more Dld endosomes are Notch^hi^, thereby supporting the notion that Dld endosomes likely contain both the ligand and receptor and hence can be considered as Notch signaling endosomes.

### Par-3 and dynein are essential for polarized segregation of Notch signaling endosomes

Par-3, an evolutionarily conserved cell polarity regulator that is asymmetrically distributed on the cell cortex ^32^, plays a critical role in establishing Notch signaling asymmetry in daughter cells of RGP ACD^4^. To determine whether Par-3 is involved in polarizing the dynamics of Notch signaling endosomes, we disrupted its activity via microinjection of a well-established morpholino (MO) antisense oligonucleotide^4,25,33^. Imaging of internalized Dld uncovered that, while Notch signaling endosomes underwent largely normal sub-cortical association and congregation toward the future cleavage plane, their final asymmetric segregation into the posterior daughter was significantly disrupted in *par-3*-deficient embryos (Figure 3b, 3e-f, Supplementary Video 9). These data indicate that Par-3 is essential for polarized segregation of Notch signaling endosomes during RGP ACD.

**Figure 3.**
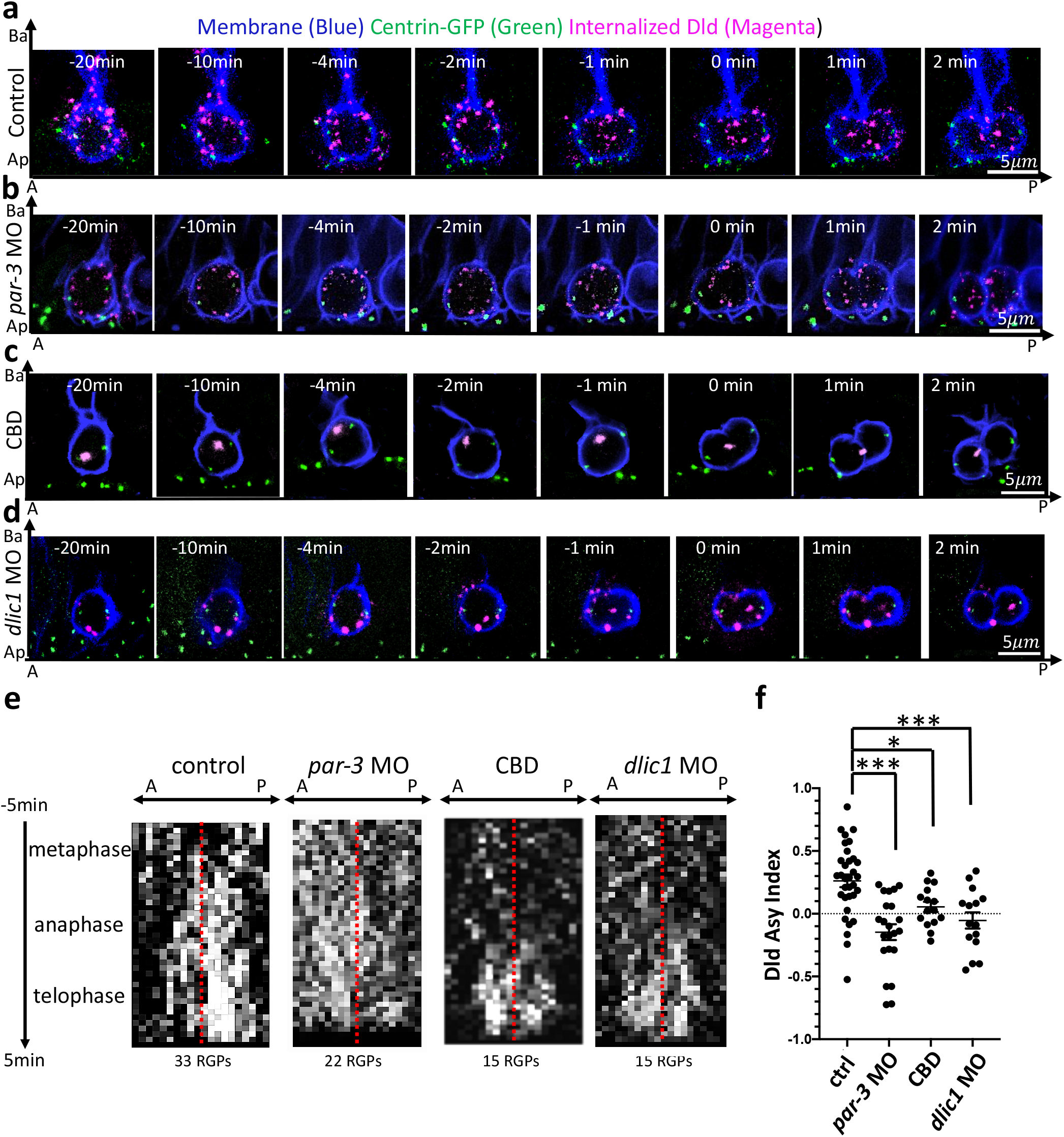
Dld endosome asymmetry is dependent on Par-3 and the dynein motor machinery. **a-d**, Time-lapse sequence of images showing Dld endosome dynamics in mitotic RGPs from 28 hpf control (a) and embryos deficient for *par-3* activity (b), treated with the dynein inhibitor ciliobrevin D (CBD)(c), deficient for dynein intermediate light chain 1 (*dlic1*)(d). Centrosomes are labeled with Centrin-GFP and membrane is labeled with Myr-TdTomato reporter. **e**, kymograph of horizontal projection of (a-d) showing distribution of all tracked Dld endosomes along the anterior-posterior (A-P) axis (x) over time (y). The red line delineates center point of the axis defined by two centrosomes. **f**, Scatter plot showing asymmetry indices in telophase RGPs. 33 control RGPs were from 25 embryos of 8 repeat experiments, 22 *par-3* MO RGPs were from 9 embryos of 6 repeat experiments, 15 CBD-treated RGPs were from 7 embryos of 4 repeat experiments, and 15 *dlic1* MO RGPs were from 8 embryos of 4 repeat experiments. *P*< 0.0001 ctrl vs *par-3* MO, t=4.706, df=53), P=0.0129 (ctrl vs CBD, t=2.589, df=46), P=0.0019 (ctrl vs *dlic1* MO, t=3.303, df=46), unpaired two tailed t-test.

Previous studies in *Drosophila* SOPs have uncovered a critical role of plus-end kinesin motors in the initial targeting of Notch signaling endosomes toward the cleavage plane^2^. However, motor involvement in the later asymmetric segregation step is not known. In dividing zebrafish RGPs, we observed directed movements of Notch signaling endosomes toward the posterior, which implies that the final polarization process might also be motor dependent. We therefore examined whether minus-end dynein motors might play a role. The pharmacological dynein inhibitor ciliobrevin D (CBD) was applied to zebrafish embryos at a concentration that still enabled RGP cell division. In CBD-treated RGPs, we observed that the Notch signaling endosomes were much larger in appearance as if they “collided” into one another. These “enlarged” endosomes remained at the cleavage furrow throughout the division (Figure 3c, 3e-f, Supplementary Video 10).

We next sought to genetically test the involvement of dynein. Since the Dynein Light Intermediate Chain 2 (DLIC2) has been reported to interact with Par-3 in cultured NIH 3T3 cells^34^, we asked whether genetic perturbation of *dlic* activity would disrupt polarized Notch signaling endosome dynamics in zebrafish RGPs. Among all the cytoplasmic dynein subunits, DLIC is the least well understood. It contains a RAS-like domain that interacts with the Dynein Heavy Chain and a C-terminal domain contacting activating adaptors and in some cases the cargo^35^. Invertebrates only express a single *dlic* gene^36,37^. Vertebrates have evolved to express two *dlic* genes (*dlic1* and *dlic2*), which may define tissue-or cell type-specific dynein complexes^38^. Using established MOs^39^ that target *dlic1* and *dlic2* in zebrafish and an anti-Dlic antibody^40^ (Extended Data Fig. 6), we found that *dlic1* is the major isoform detected in mitotic RGPs and its activity is essential for asymmetric posterior targeting of Notch signaling endosomes (Figure 3d, 3e-f). Together, combined pharmacological and genetic perturbation of dynein function, while not affecting the congregation of Notch signaling endosomes toward the cleavage plane, selectively disrupts their asymmetric segregation into the posterior daughter during RGP ACD.

### Cytoplasmic Par-3 and dynein decorate Notch signaling endosomes

Now that we have established an essential role of Par-3 and the dynein motor in polarizing the distribution of Notch signaling endosomes during RGP ACD, we next asked how they might perform such function. In both invertebrates and vertebrates, Par-3 protein displays prominent cortical asymmetry and is widely considered to act at the cell cortex/membrane during ACD^25,32,41,42^. To understand how Par-3 cortical asymmetry relates to polarized Notch signaling endosome dynamics, we performed *in vivo* time-lapse imaging to simultaneously track the dynamics of Par-3 and internalized Dld in dividing RGPs (Figure 4a, Supplementary Video 11). Par-3-GFP, which is shown fully functional through rescue experiments^33^, was used as a live reporter. We found that, during metaphase, Par-3 was prominently localized to the apical cortex. Starting at around anaphase, however, weak but discernable cytoplasmic Par-3 was detected preferentially in the posterior half (Figure 4a, arrows). Some GFP signals were found to be in close proximity to Dld endosomes. By the end of telophase, Par-3 again was cortical and became enriched in the posterior daughter, which also had more Dld endosomes. Quantification of Par-3-GFP (both cytosolic and membrane) and Dld endosome distribution in 22 telophase RGPs exhibiting Par-3 posterior enrichment showed a strong positive correlation (Extended Data Fig. 7).

**Figure 4.**
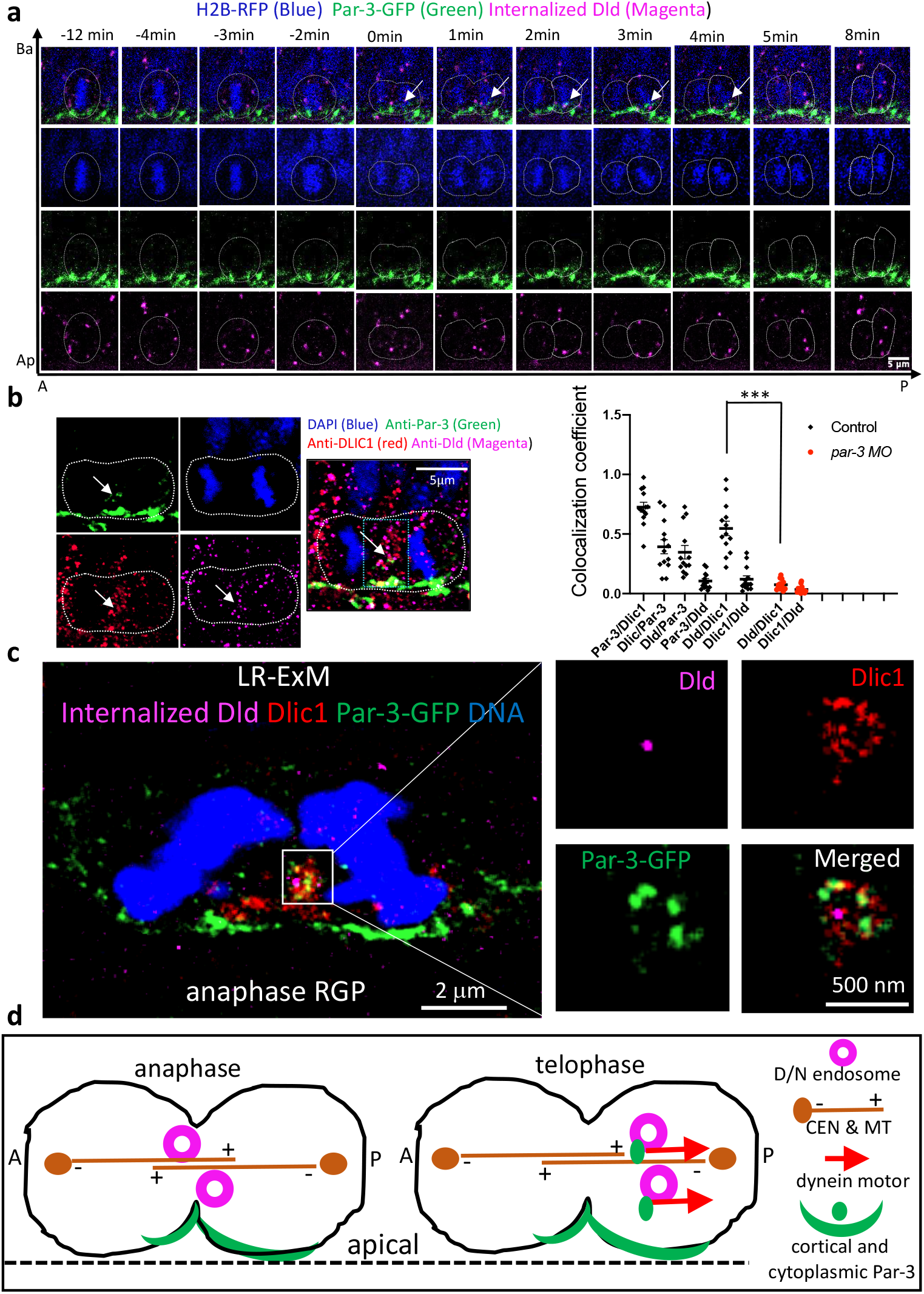
Cytosolic Par-3 together with Dlic1 decorates Dld endosomes. **a**, Time-lapse sequence of images showing the dynamics of internalized Dld and Par-3-GFP in mitotic RGPs. Arrows point to the cytosolic Par-3-GFP in close proximity to internalized Dld. DNA is marked by H2B-RFP (pseudo-colored in blue). **b**, Immunofluorescent (IF) labeling with anti-Par-3, anti-Dlic1, and anti-Dld in anaphase RGP (left). (right) Quantification of co-localization coefficients (See Methods) in control vs. *par-3* deficient RGPs. The region used for quantification is indicated by the green box (60×100 pixel = 7×12*μ*m) in the merged IF image. For instance, Par-3/Dlic1 indicates the percentage of co-localized Par-3 and Dlic1 among total labeled Par-3. Par-3 IF was diminished in *par-3* MO-injected embryos (See Extended Data Fig. 9). Dld/Dlic1 co-localization was significantly reduced in the absence of *par-3*. *P*< 0.001 (*t* = 7.726, and *df* = 24), unpaired two-tailed t-test, n=13 RGPs for each genotype from 3 repeat experiments. Mean with S.E.M was shown for each group.**c**, Label Retention-Expansion Microscopy (LR-ExM) of anaphase RGP from 28 hpf embryo injected with *par-3*-GFP mRNA at 16-cell stage and anti-Dld-Atto647 antibody at 26 hpf. The sample is expanded ~4 fold. Both Par-3 and Dlic1 decorate a ring-like structure containing Dld with approximate real biological size of < 500 nm (the average size of an endosome). The left whole cell image is max projection of 10 z-planes; the right enlarged views are max projection of 4 z-planes for visualizing the ring-like structure of endosome. 14 RGPs from 4 repeat experiments were analyzed for patterns of protein co-localization (See Extended Data Fig. 9). **d**, model summarizing the finding. D/N, delta and Notch; CEN, centrosome; MT, microtubule.

Prior studies are focused on the membrane-associated Par-3, which is much more abundant, but we were intrigued by the cytosolic Par-3 and its apparent proximity to Dld endosomes. The cytosolic presence of Par-3 protein was further verified using an anti-Par-3 antibody, the specificity of which was validated using the *par-3* MO (Extended Data Fig. 8). Co-immunofluorescence (IF) labeling of Par-3, Dlic1, and Dld was then analyzed (Figure 4b), using a method that is shown to work well for quantifying co-localization in the context of endosomes^2,43^. It was found that in the cytoplasmic area in between two nuclei of the anaphase/telophase RGPs, ~60% of total Par-3 immunoreactivity co-localized with Dlic1, whereas about 30% of total Dlic1 immunoreactivity co-localized with Par-3. This difference was expected given the broad role of Dlic1 in other dynein-mediated processes. It was further observed that the co-localization of Dld with Dlic1 was significantly decreased in *par-3* deficient embryos, suggesting an essential role of Par-3 in linking Dlic1 (in turn the dynein motor) to Notch signaling endosomes.

To visualize at high resolution the cytoplasmic Par-3 in relationship to that of Dld endosomes and Dlic1, we applied label-retention expansion microscopy (LR-ExM). ExM physically expands biological samples to enable nanoscale super-resolution via diffraction-limited confocal microscopy^44,45^. LR-ExM further overcomes the limitation of signal loss associated with traditional ExM and enables high-efficiency protein labeling using an engineered set of trifunctional anchors^46^. 24-28 hpf embryos expressing Par-3-GFP and with internalized Dld were processed for immunolabeling of GFP and Dlic1 followed by LR-ExM.

In addition to the cortex, Par-3-GFP was detected in the cytosol in close proximity to Dlic1 and Dld (Figure 4c). The ring-shaped distribution patterns of Dlic1 and Par-3 together with the diameter of the ring suggest that they decorate Dld endosomes. We also performed LR-ExM by immuno-labeling of endogenous Par-3. The results showed similar co-localization of endogenous Par-3 and Dlic1 on Dld endosomes (Extended Data Fig. 9).

## Discussion

The evolutionarily conserved polarity regulator Par-3 is required for differential activity of Notch signaling in daughter cells of asymmetrically dividing RGPs^4^. Using *in vivo* time-lapse imaging and expansion microscopy, in combination with molecular genetics and pharmacological approaches, we have uncovered a previously unknown mechanism: Par-3 together with the dynein motor are essential for polarized transport of Notch signaling endosomes, and they do so by directly associating with endosomes in the cytoplasm. This study, together with previous work^2^, reveals an evolutionarily conserved role of Par-3 in polarizing Notch signaling endosomes. However, it appears to do so through distinct processes: in *Drosophila* SOPs, Par-3 regulates spindle asymmetry^2^; in zebrafish RGPs, Par-3 engages dynein in direct transport of endosomes.

During ACDs studied across different metazoan species, Par-3 displays prominent localization at distinct membrane compartments of mitotic progenitors^25,32,41,42^. This membrane-associated form of Par-3 has received much attention as key to establishing polarity^27^. Cytoplasmic Par-3, released from the membrane via phosphorylation by Par-1^47–50^, has been considered a largely non-functional form in the context of polarity regulation. Being able to visualize Par-3 in the cytoplasm in association with the dynein motor and Notch signaling endosomes as we have done in this study, for the first time to our knowledge, reveals a direct role of Par-3 in actively localizing cytosolic determinants. Cytoplasmic Par-3 has been associated with adverse cancer prognosis in clinical settings^51^, further suggesting that this form of Par-3, and its dynamic relationship with the membrane-associated oligomeric ensemble, deserve more attention in future studies.

## Acknowledgments

We thank Dr. Mahendra Wagle for technical training on brain ventricle-targeted electroporation, Michael Munchua and Vivian Yuan for excellent animal care, Drs. Arnold Kriegstein, Lily Y. Jan, Yuh Nung Jan, Bingwei Lu, and Guo lab members for helpful discussion, DeLaine Larsen, Kari Herrington and UCSF Nikon imaging center for assistance with imaging and data analysis. This project was supported by NIH R01NS095734 to S.G., the UCSF Mary Anne Koda-Kimble Seed Award for Innovation (to X.Z.), Fudan Bio-elite undergraduate program (to K.T.), NIH R01GM124334 and R01GM131641 (to B.H.), and NIH K99GM126136/R00GM126136 (to X.S.). B.H. is a Chan Zuckerberg Biohub Investigator.

## Author contributions

X.Z. and S.G. designed the experiments and interpreted the results. X.Z., and K.T. performed the experiments, X. S., I.S., and B.H. assisted with critical steps in establishing LR-ExM on *in vivo* zebrafish samples, X. C., B.Y. and B.H. assisted with critical steps in image analysis. X.Z. and S.G. wrote the manuscript, with the input from all authors.

## Materials & Correspondence

Correspondence and material requests should be addressed to S.G.

## METHODs

### Zebrafish strains and maintenance

Wild-type embryos were obtained from natural spawning of AB adults, staged and maintained according to established protocols^52^. Embryos were raised at 28.5 °C in 0.3x Danieau’s embryo medium (30x Danieau’s embryo medium contains 1740 mM NaCl, 21 mM KCl, 12 mM MgSO_4_•7H_2_O, 18 mM Ca(NO_3_)_2_, 150 mM HEPES buffer). Embryonic ages were described as hours post-fertilization (hpf). To prevent pigment formation, 0.003% Phenylthiourea (PTU) was added into the medium at 24 hpf. The following zebrafish mutants and transgenic lines were used: *mib^ta52b^*^29^, *Tg [ef1a:Myr-Tdtomato]* and *Tg[ef1a:H2B-RFP]*^53^, and *Tg [her4.1-RFP]*^31^. All animal experiments were approved by the Institutional Animal Care and Use Committee (IACUC) at the University of California, San Francisco, USA.

### Morpholinos

Knockdown experiments were carried out using previously characterized translational blocking antisense morpholino oligonucleotides (MOs): *par-3* MO (5’-TCA AAG GCT CCC GTG CTC TGG TGT C-3’)^4,25,33,54^; *dlic1* MO 5’-GTG TAT TTC TGC CCG TCG TCG CCA T-3’) and *dlic2* MO (5’-TTC TTC TCT AAA ACGGGA GCC ATC T-3’)^39^. Standard control MO 5’-CCT CTT ACC TCA GTT ACA ATT TAT A-3’) was used as injection controls (Gene Tools). All MOs were stored at 300 mM in distilled water. For microinjection, ~4 nl of the diluted MO at 100 mM in the injection mixture containing 0.05% phenyl red (corresponding to 4 ng of MOs) was injected into the yolk of 1-4 cell stage embryos.

### Pharmacology

Zebrafish embryos were treated with the dynein inhibitor ciliobrevin D (CBD)(Calbiochem via Sigma-Aldrich, 250401-10MG)(2.5 μM in 0.1% DMSO, 0.3x Danieau’s buffer) from 18 hpf to 24 hpf. Vehicle (DMSO)-treated control and CBD-treated embryos were then embedded in 1.2% low-melt point agarose in the Danieau medium supplemented with 2.5 μM ciliobrevin D and 0.003% Tricaine and subjected to Dld antibody uptake assay and *in vivo* time-lapse imaging.

### DNA Plasmids

Plasmid DNAs (*pCS2-H2B-mRFP; pCS2-mib-GFP; Pef1a-gal4; Puas-E1b-EGFP)* were prepared as previously described^4,55^. *pCS2-par-3-GFP* plasmid was a gift from Dr. J. von Trotha^56^. *pCS2-GFP-centrin* plasmid was a gift from Dr. W. A. Harris^57^. For *mib-GFP*, GFP is at the 3’ end of Mib protein. For *par3-GFP*, GFP is at the 3’ end of Par-3 protein. For *GFP-centrin*, GFP is at the 5’ end of Centrin.

### mRNA synthesis and microinjection

Plasmids (*pCS2-H2B-mRFP; pCS2-mib-GFP; pCS2-par-3-GFP; pCS2-GFP-centrin*) were linearized by the restriction enzyme Not I digestion. NotI-linearized plasmids were purified (QIAquick Gel Extraction Kit) and the 5’-capped mRNAs were synthesized using SP6 mMessenger mMachine kit (Ambion). For *GFP-centrin*, *H2B-mRFP*, and *mib-GFP* mRNA injection, 4nl mRNAs at 0.2 *μ*g/*μ*l were mixed with equal volume of injection buffer containing 0.05% phenyl red and injected into the yolk of a 1-4 cell stage embryos. For *par-3-GFP* mRNA injection, the mRNAs were injected into single cells at the 16–32 cell stage to obtain mosaic expression. All injections were done with an injector (WPI PV830 Pneumatic Pico Pump) and a micromanipulator (Narishige, Tokyo, Japan).

### Anti-Dld antibody uptake assay

Anti-Mouse-IgG-Atto647N (Sigma-Aldrich) was used for labeling the mouse monoclonal anti-Dld antibody (Abcam ab73331). For antibody conjugation, 1 μl of anti-Dld antibody (0.5mg/ml) was mixed with 2.5μl Anti-Mouse-IgG-Atto647N antibody (1mg/ml) and incubated at room temperature for 30 minutes or on ice for 2-3 hours. After incubation, 2.5 μl blocking buffer (10 mg/ml mouse IgG with 5 mM Azide) and 0.5 μl 0.5% Phenol red (Sigma P0290) were added for blocking the unconjugated Anti-Mouse-IgG-Atto647N and vortexed thoroughly. Mixtures without anti-Dld antibody were used as control. Before microinjection, 24-26 hpf embryos were anaesthetized in the Danieau medium supplemented with 0.003% Tricaine followed by embedding in 1.2% low-melt point agarose. 10 nl of labelled Dld antibody was injected into the hindbrain ventricle. The phenol red indicator serves to show the diffusion of antibody mixture into the forebrain ventricle. The injected embryos were then released from agarose and cultured in the Danieau medium for 2 hours before imaging.

### Brain ventricle-targeted electroporation of plasmid DNAs

*Pef1a-gal4* and *Puas-E1b-EGFP* plasmids were mixed at 500 ng/μl for each, and about 20 nl mixture was microinjected into the hindbrain ventricles of 20-22 hpf *Tg [her4.1-RFP]* embryos embedded in 1.2% low-melt point agarose. The electroporation setting and procedures were according to previously established protocols^55^. Electroporated embryos were released from agarose and transferred to a fresh dish of embryonic medium containing 0.003% PTU and incubated at 28.5°C. Electroporated embryos were checked under a fluorescent stereo microscope after 6-8 hours. The embryos with sparsely GFP-labeled RGPs were subjected to the Dld antibody uptake assay followed by *in vivo* time-lapse imaging.

### Antibodies, western blotting, and immunocytochemistry

Primary antibodies used in this study were: mouse anti-Dld (Abcam ab73331; RRID:AB_1268496; lot GR115501-3, 1:200 dilution for immunostaining)^4^, chicken anti-GFP (Abcam ab13970; RRID:AB_300798, lot GR3190550-20, 1:500 dilution for immunostaining), rabbit anti-Par-3 (Millipore 07-330; RRID:AB_2101325; lot 3322358, 1:500 for immunostaining)(validated in this study), guinea pig anti-DLIC-Cter (a gift from D. T. Uemura, 1:100 for immunostaining and 1:500 for western blotting, validated in this study)^40^, rabbit anti-α-Tubulin (Santa Cruz SC-12462; RRID:AB_2241125, lot A2907, 1:1000 for western blotting).

Secondary antibodies used for immunofluorescent labeling were: Alexa®-conjugated goat anti-rabbit (Alexa 568, Invitrogen A11011, RRID. AB_143157, lot 792518), goat anti-chicken (Alexa 488, Invitrogen A11039, RRID:AB_142924, lot 2020124), goat anti-mouse (Alexa 488, Invitrogen A11002, RRID:AB_2534070, lot 1786359), goat anti-guinea pig (Alexa 488, Invitrogen A11073, RRID:AB_2534117, lot 46214A) or donkey anti-guinea pig (Alexa 647, Jackson Labs 706-605-148, RRID:AB_2340476, lot 102649-478). All were used at 1:1000 dilution.

Secondary antibodies used for western blotting were: Amersham ECL Donkey anti-Rabbit IgG HRP (GE Healthcare NA934V, RRID:AB_772211, lot 16921443) and Rabbit anti-guinea pig IgG (H+L) HRP (Sigma-Aldrich A5545, RRID:AB_258247). All were used at the 1:2000 dilution.

Secondary antibodies used for tri-functional linker conjugation in LR-ExM were: Goat anti-Guinea Pig IgG (H+L) unconjugated secondary antibody (Invitrogen A18771, RRID:AB_2535548), Goat anti-Rabbit IgG (H+L) Cross-Adsorbed unconjugated secondary antibody (Invitrogen 31212, RRID:AB_228335) and Goat anti-Chicken IgY (H+L) unconjugated secondary antibody (Invitrogen A16056, RRID:AB_2534729).

For Western blotting, lysates from 15–20 28 hpf embryos were homogenized in 80 μl SDS sample buffer; 15 μl lysate was used for SDS-PAGE (BioRad). Proteins were transferred to HybondTM nitrocellulose membrane by a semidry blotting technique with Turbo-transblotting cell (Bio-Rad) and detected with appropriate antibodies as previously described^58^. After the horseradish peroxidase-conjugated secondary antibody incubation, the samples were visualized using the SuperSignal™ West Dura Extended Duration Substrate (Thermo Scientific™) with the LI-COR Western blotting detection system (LI-COR Biosciences).

For the preparation of cryosections, 28 hpf embryos were fixed overnight at 4 °C in phosphate-buffered saline (PBS) buffer with 4% paraformaldehyde. Fixed embryos were washed and incubated in PBS buffer containing 30% sucrose overnight at 4 °C. Embryos were then transferred to plastic molds, with the sucrose buffer removed and OCT (Tissue-tek) added. After orienting the embryos to proper positions in the mold, the block was frozen on dry ice. Blocks can be stored at −80 °C up to several months. Frozen blocks were then cut into 20 μm sections on a cryostat (Leica) and mounted on Superfrost Plus slides (Fisher Scientific). The slides were dried at room temperature for 2-3 hours and then stored at −80 °C until use.

For immunocytochemistry, samples were first washed and preincubated in phosphate-buffered saline containing 0.1% Tween 20 or 0.25% Triton X-100 (PBS-T, pH 7.4) with 1% dimethyl sulfoxide (DMSO) and 5% natural goat serum at 4 °C overnight or longer. They were then incubated with primary antibodies in the preincubation solution (PBS-T with 5% natural goat serum) overnight at 4 °C. The samples were then washed thoroughly with PBS-T 5 times × 10 minutes each time, followed by incubation in Alexa®-conjugated goat anti-rabbit (Alexa 568), goat anti-chicken (Alexa 488), goat anti-mouse (Alexa 488) or goat anti-guinea pig (Alexa 647) secondary antibodies (diluted 1:1000) in the preincubation solution for over 2 hours at room temperature or overnight at 4 °C. The samples were washed with PBS-T twice for 10 minutes each, three times with PBS for 10 minutes each, and once with 50% glycerol in PBS for 1 hour, followed by infiltration overnight in 80% glycerol/PBS before imaging. Imaging was done using the confocal microscope (Nikon CSU-W1 Spinning Disk/confocal microscopy) with 100X oil immersion objective. The z-step of the imaging stack is 0.26 μm.

### Time-lapse *in vivo* imaging

Time-lapse *in vivo* imaging was done using the confocal microscope (Nikon CSU-W1 Spinning Disk/High Speed Widefield confocal microscopy) with 40X water immersion objective. Embryos were mounted with 1.2% low-melting-point agarose (0.3x Danieau medium and 0.003% tricaine) in glass bottom culture dishes (MatTek 35mm) with the dorsal forebrain facing the coverslip.

For *in vivo* time-lapse imaging of internalized Dld, Mib-GFP, or Par-3-GFP in dividing RGPs, z-stacks ranging from 20 to 80 were acquired consecutively with a z-step at 1 μm for each embryonic forebrain region. The scanning interval ranged from 12 sec to 10 min, and imaging duration spanned from 20 min to 12 hr. For long-term imaging, embryos were placed on a temperature-controlled stage set at 28.5°C. For imaging Notch activity in paired daughter cells using *Tg[her4.1-RFP]* embryos, a GFP reporter plasmid was electroporated into the hindbrain region to label individual RGPs as this transgenic line was reported to better recapitulate Notch activity in the hindbrain than in the forebrain^31^. Data presented in figure panels corresponded to max-intensity projections of 5-7 z-planes with 1 μm z-step size, approximately covering the size of RGP.

### Expansion Microscopy

Expansion microscopy was performed on cryosections of 28 hpf embryonic forebrain. Chicken anti-GFP antibody (for detecting Par-3-GFP) or rabbit anti-Par-3 antibody (for detecting endogenous Par-3) and Guinea Pig anti-Dlic-Cter antibody were used, in conjunction with visualizing Dld (either internalized Atto-labeled Dld or endogenous Dld labeled with mouse-anti-Dld antibody). Buffers were prepared as previously described with modifications needed for processing *in vivo* tissue samples^46^. The forebrain sections were blocked in blocking buffer, phosphate-buffered saline containing 0.1% Triton X-100 (PBS-T, pH 7.4) with 5% natural goat serum overnight at 4 °C. The slides were then incubated with primary antibodies in the preincubation buffer overnight at 4 °C as described in the immunocytochemistry section above. After washing off primary antibodies, tissue sections were incubated with trifunctional linkers (NHS-MA-Biotin conjugated anti-Chicken (or anti-Rabbit) IgG (for visualizing Par-3), and NHS-MA-DIG conjugated anti-Guinea Pig IgG (for visualizing Dlic1) (200 μg/ml stock, dilute 1:100 before use) in the preincubation buffer overnight at 4 °C in dark. After washes in PBS 4 times (5 min each) in dark, freshly prepared 40 μl gelation solution was added on each section to cover the whole tissue sample. The gelation solution was prepared by deoxygenizing the gel monomer solution using a vacuum pump for over 15 minutes before adding TEMED and APS, to enhance the effects of trifunctional linkers. The gelation solution-covered samples, protected from light, were incubated in a humidity chamber and allowed to undergo gelation at 37 °C for 1 hour. The gelated samples were incubated in the digestion buffer (8 units/ml Proteinase K in 50 mM Tris pH 8.0, 1 mM EDTA, 0.5% Triton X-100) on the slides 4 h at 37 °C or overnight at room temperature. At least 10-fold excess volume of digestion buffer was used. Sufficient digestion enabled sections embedded in gels to slide off the glass surface. The gels were washed with excess volume of 150 mM NaCl in 6-well plates (black-walled plates, CellVis P06-1.5H-N), at least 5 ml in each well for 4 times, 20-30 min each time. After washing off the digestion buffer, the gel samples were incubated in the staining buffer (10 mM HEPES, 150 mM NaCl, pH=7.5) with 3-5 *μ* M Alexa Fluor® 488-Streptavidin, 3-5 *μ* M goat anti-Digoxigenin/Digoxin Dylight® 594, anti-mouse Atto647N (1:500), and DAPI (1:1000) for 24 hour at 4 °C in dark. Finally, the gels were washed 4 times with milli-Q water (30 min for each wash) at 4 °C in dark. The gel expanded approximately four times of the original sample size after washing and was ready for imaging in the well under the confocal microscope (Nikon CSU-W1 Spinning Disk/High Speed Widefield confocal microscopy) with 60X water immersion objective. The excess water in the well was removed in order to keep the gel embedded samples adhered to the glass dish bottom. For immobilization of the gel, 2% low melt agarose was added onto the edge of the gel embedded samples before imaging. The scanning z-step is 0.26 μm.

### Image analyses

All the confocal imaging stacks were processed using Micro-Manager and Fiji. For generating kymographs of internalized Dld in dividing RGPs, Max Intensity Projection (MIP) was applied to the 3D image stacks to cover the entire RGP. Each RGP was manually segmented according to the cell membrane labeling. Each single Dld endosome was located from all frames using a Gaussian fitting algorithm assisted with the ImageJ embedded plugin TrackMate^59^. The two labeled centrosomes were used to define the anterior-posterior axis: the anterior centrosome was given the coordinate of 0 and the posterior centrosome 1. Each Dld endosome was then projected onto this axis to obtain its relative distance (value between 0 and 1). The relative distances of all tracked Dld endosomes are then used to generate the kymograph, where the grayscale value of each pixel indicates the probability of Dld endosomes appearing at the corresponding location at a given time.

Movies for each RGP were registered spatiotemporally: The spatial registration was done by adopting the center point between two centrosomes as the center of dividing RGPs. The temporal registration was done by adopting the onset of anaphase to be t=0. The onset of anaphase was recognized as first appearance of the cleavage furrow, which was verified in a set of embryos with double labeling of cell membrane and nucleus. For *par-3* MO, *dlic* MO, or CBD-treated embryos, since their cell cycle lengths sometimes appeared altered, we normalized these to WT cell cycle by assigning the onset of telophase (i.e. with two newly formed daughter cells) as t= −2 min.

### Quantification and statistical analysis

The number of times each experiment was repeated was provided in the figures or figure legends. For live imaging, one or multiple RGPs were analyzed from each embryo, depending on the number of mitotic RGPs that were present in each image stack. For immunocytochemistry experiments, multiple sections from individual brains were analyzed. No Statistical methods were used to predetermine sample size. Sample size was determined to be adequate on the basis of the magnitude and consistency of measurable differences between groups. No randomization of samples was performed. Embryos used in the analyses were age-matched between control and experimental conditions, and sex cannot be discerned at these embryonic stages. Investigators were not blinded to chemically or genetically perturbed conditions during experiments. Data are quantitatively analyzed.

Statistical analyses were carried out using Prism 8 version 8.4.2: the mean value with standard error of the mean (S.E.M.) was labeled in the graphs. The two-tailed *t*-test was used to asses significance. To compare the proportions of each cell division orientation (Extended Data Fig. 1), normal (z) test for proportions (implemented by Python’s statsmodels package^60^) was used.

### Measurement of asymmetry index

The total fluorescence intensity of internalized Dld (or Mib-GFP, Par-3-GFP) in paired daughter cells immediately after abscission (i.e. at telophase of mother RGP division) was measured by FIJI. To quantitatively describe the distribution, normalized ratio of fluorescence between the two newly formed daughter cells was calculated as follows:

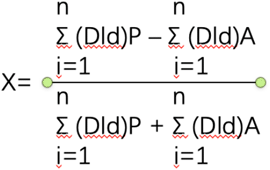

Σ(Dld)P means total intensity in the posterior daughter cell, and Σ(Dld)A means total intensity in the anterior daughter cell. And “0” indicates perfect symmetry and “1” or “-1” indicates absolute asymmetry (posterior or anterior, respectively). For filtering out potential noise, we defined asymmetry when Σ(Dld)P is 50% more or less than Σ(Dld)A, as has been previously used^6^. It means that when X ≥ 0.2, Dld endosomes (or Mib-GFP, Par-3-GFP) are considered asymmetric with more in the posterior daughter, and when X ≤ −0.2, they are considered asymmetric with more in the anterior. The asymmetry index for Par3-GFP included both membrane and cytoplasmic fluorescence.

### Co-localization

To measure the co-localization of Dlic1, Par-3 (or Par-3-GFP), and Dld (either internalized or immuno-stained) in the cytosol of dividing RGPs, we adopted the methods as previously described for measuring co-localization in the context of endosomes^2,43^. Immuno-stained RGPs were segmented and max intensity projections of 20 consecutive z-planes (with 0.26 *μ*m z-step size) were generated, which typically covers the cytoplasmic area of a mitotic RGP (~5 *μ*m). The central area of 60 x 100 pixels (1 pixel = 0.126 μm) of a mitotic RGP (i.e. the cytoplasmic area between nuclei) was then cropped for co-localization analysis using the JACop plugin in Image J. The colocalization between each fluorescent channel (i.e. the proportion of each fluorescence colocalized with another) was measured using Manders’ Coefficients^61^. We used an optimized XY block size of 2 pixels^43^. To remove potential background noise, the threshold of each channel was set in JACop using a blank area on the image (i.e. without tissue samples) as negative control. Costes’ automatic threshold was further applied for each measurement. Costes’ automatic threshold is an algorithm to identify and remove noise using scatter plotting of randomized images generated from the image under analysis^62^. It automatically quantifies co-localization without the bias of visual interpretation.

For RGPs from LR-ExM, the max intensity projections of 10 consecutive z-planes (0.26 *μ*m z-step size scanned with a 60 X water immersion objective) were generated. The cytoplasmic area of RGP between the nuclei was then selected and XY block size was defined as 1 pixel for JACop co-localization analyses.

## Extended Data Figures

**Extended Data Figure 1.**
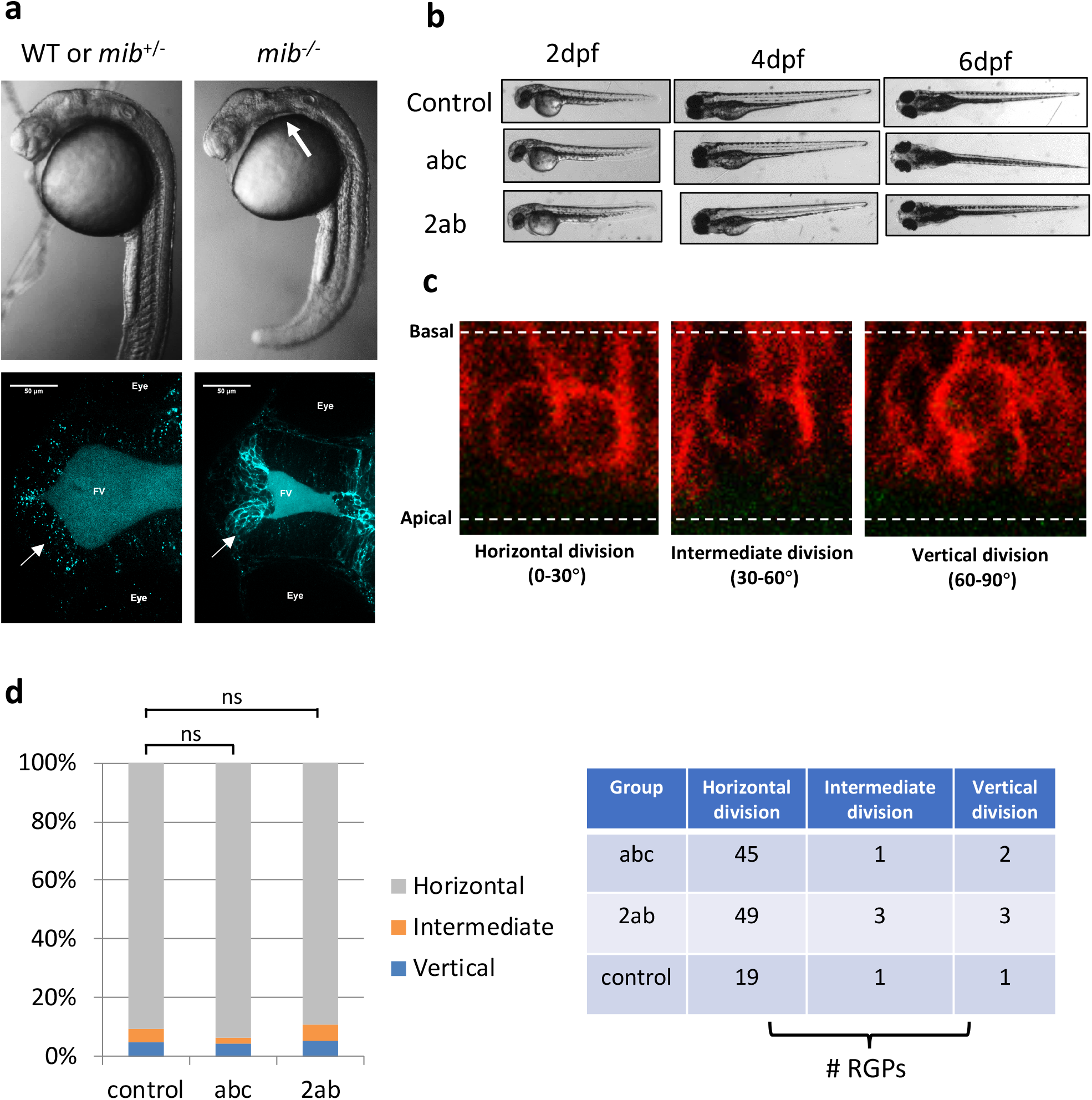
*In vivo* antibody uptake assay labels endocytosed Notch ligand Dld without affecting cell division patterns and embryonic development. **a**, morphological (top) and confocal images (bottom) showing impaired Dld internalization due to a defect in Notch ligand endocytosis in 24 hpf *mib*^−/−^ embryo compared to control sibling. Similar results were observed in 5 embryos for each condition. **b**, Morphological images showing grossly normal development of control, abc (injected with the dye-labeled secondary antibody conjugated anti-Dld), and 2ab (injected with the dye-labeled secondary antibody only) embryos at 2 dpf, 4dpf and 6 dpf (days post fertilization). Similar results were observed in 5 embryos for each condition. **c**, Confocal images showing different orientations of division in mitotic RGPs from ~28 hpf embryos (horizontal division, intermediate division and vertical division). **d**, Statistical graph showing percent division modes of RGPs from ~28 hpf embryos in control, abc, and 2ab. Normal (z) test for proportions (implemented by Python statsmodels package) shows no statistical difference between control group and any of the other two groups. Control vs abc (Proportions of Horizontal division): zscore = −0.483, *P* = 0.629; control vs abc (Proportions of Intermediate division): zscore = 0.610, *P* = 0.542; control vs abc (Proportions of Vertical division): zscore = 0.112, *P* = 0.911; control vs 2ab (Proportions of Horizontal division): zscore = 0.176, *P* = 0.860; control vs 2ab (Proportions of Intermediate division): zscore = −0.121, *P* = 0.904; control vs 2ab (Proportions of Vertical division): zscore = −0.121, *P* = 0.904.

**Extended Data Figure 2.**
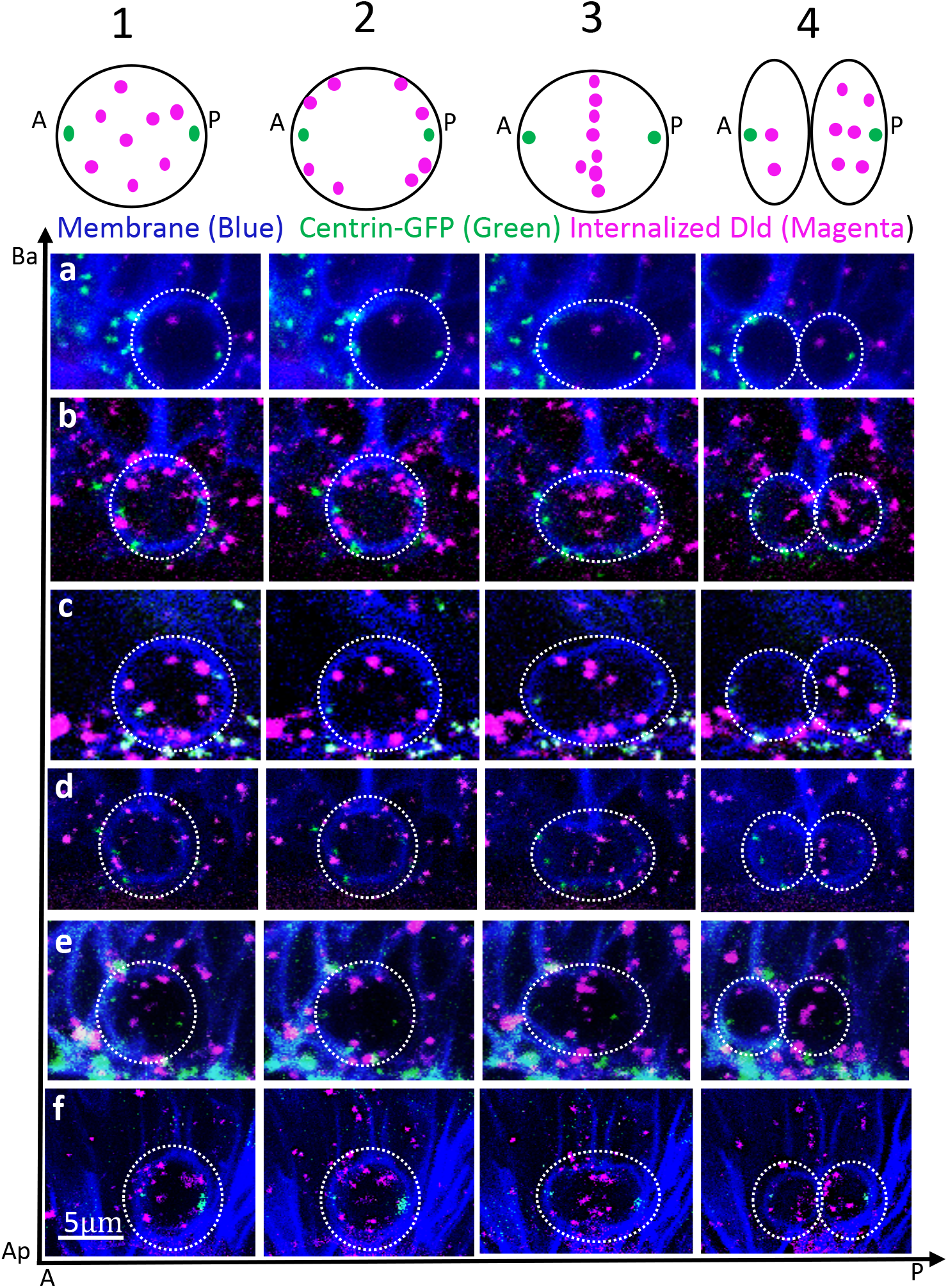
*In vivo* time-lapse sequences of images show the dynamics of Dld endosomes in 6 additional asymmetrically dividing RGPs. (Top) Schematic showing the four phases of Dld endosome dynamics. **a-f**, six mitotic RGPs. Membrane is marked with Myr-TdTomato (pseudo-colored blue). Centrosome is marked with GFP-Centrin.

**Extended Data Figure 3.**
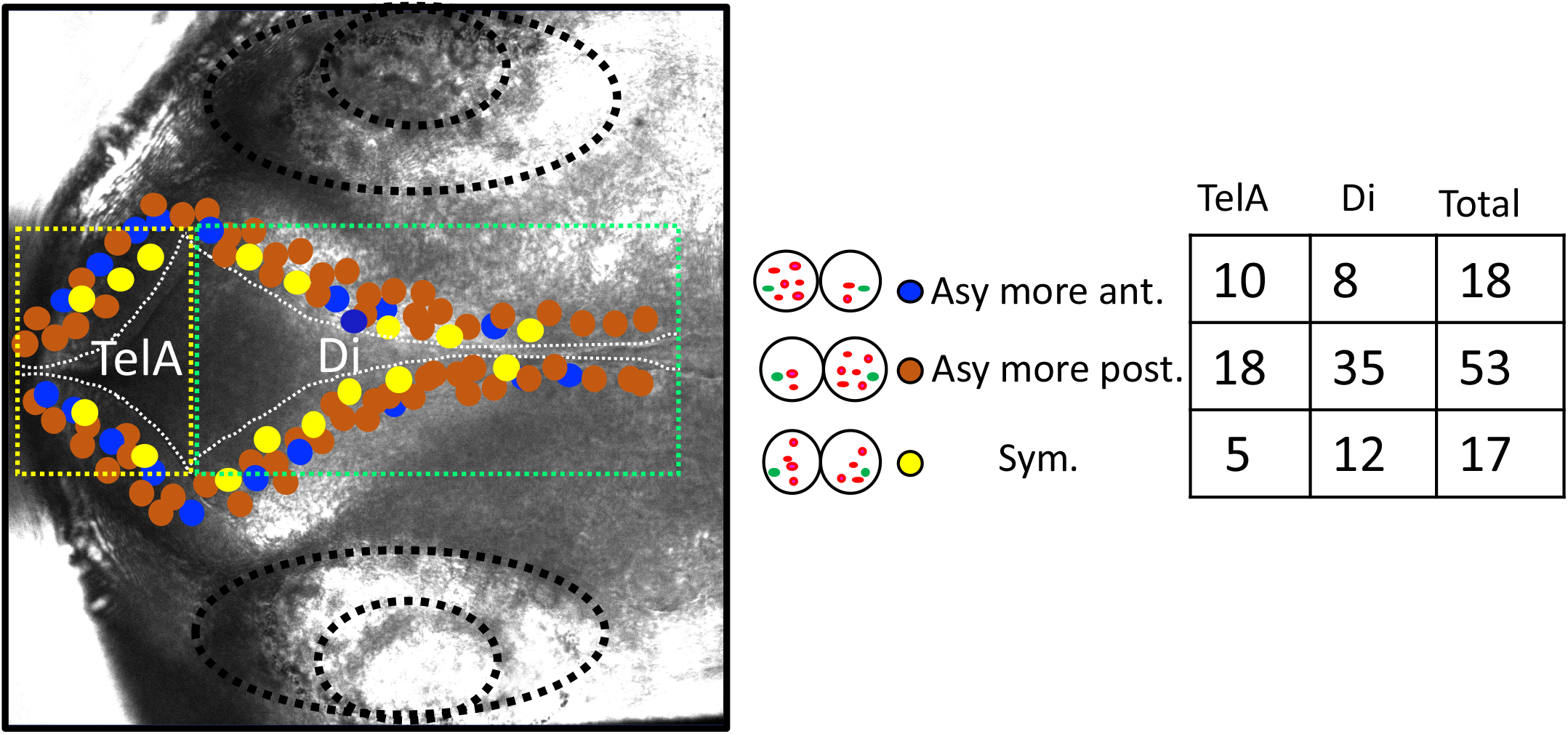
Spatial distributions of imaged telophase RGPs in the embryonic forebrain. Bright-field image of 28 hpf embryonic forebrain. The position of telophase RGPs (n=88) surrounding the forebrain ventricle was plotted. These RGPs were included in the statistics of **Figure 1d**. TelA, telencephalic anterior part of the ventricle; Di, diencephalic part of the ventricle.

**Extended Data Figure 4.**
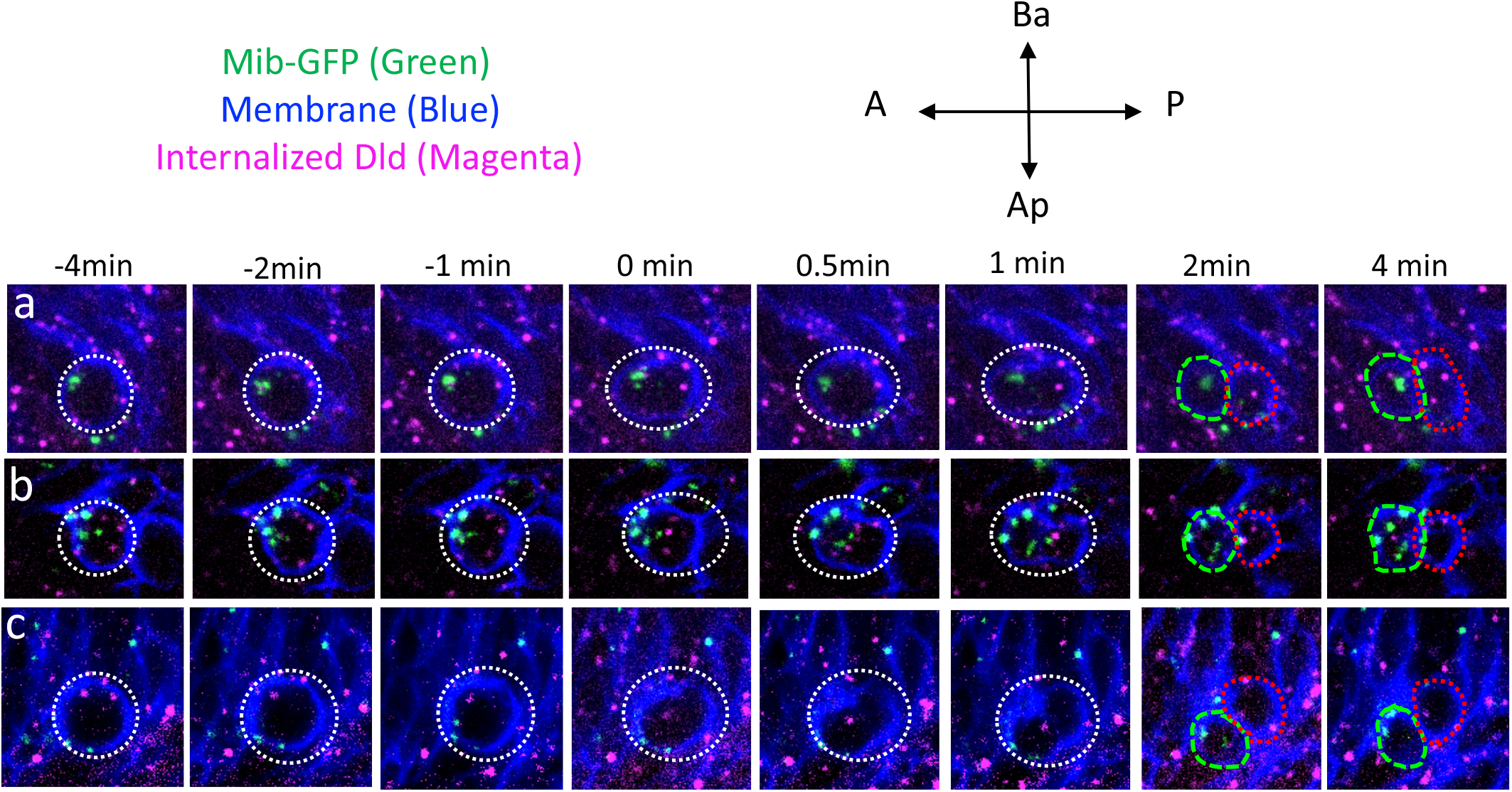
Mib and Dld endosomes preferentially segregate to different daughter cells in 3 additional asymmetrically dividing RGPs. **a-c**, *In vivo* time-lapse sequence of images showing three mitotic RGPs, with more Mib-GFP in the anterior daughter and more Dld endosomes in the posterior daughter. The membrane is marked with Myr-TdTomato (pseudo-colored blue).

**Extended Data Figure 5.**
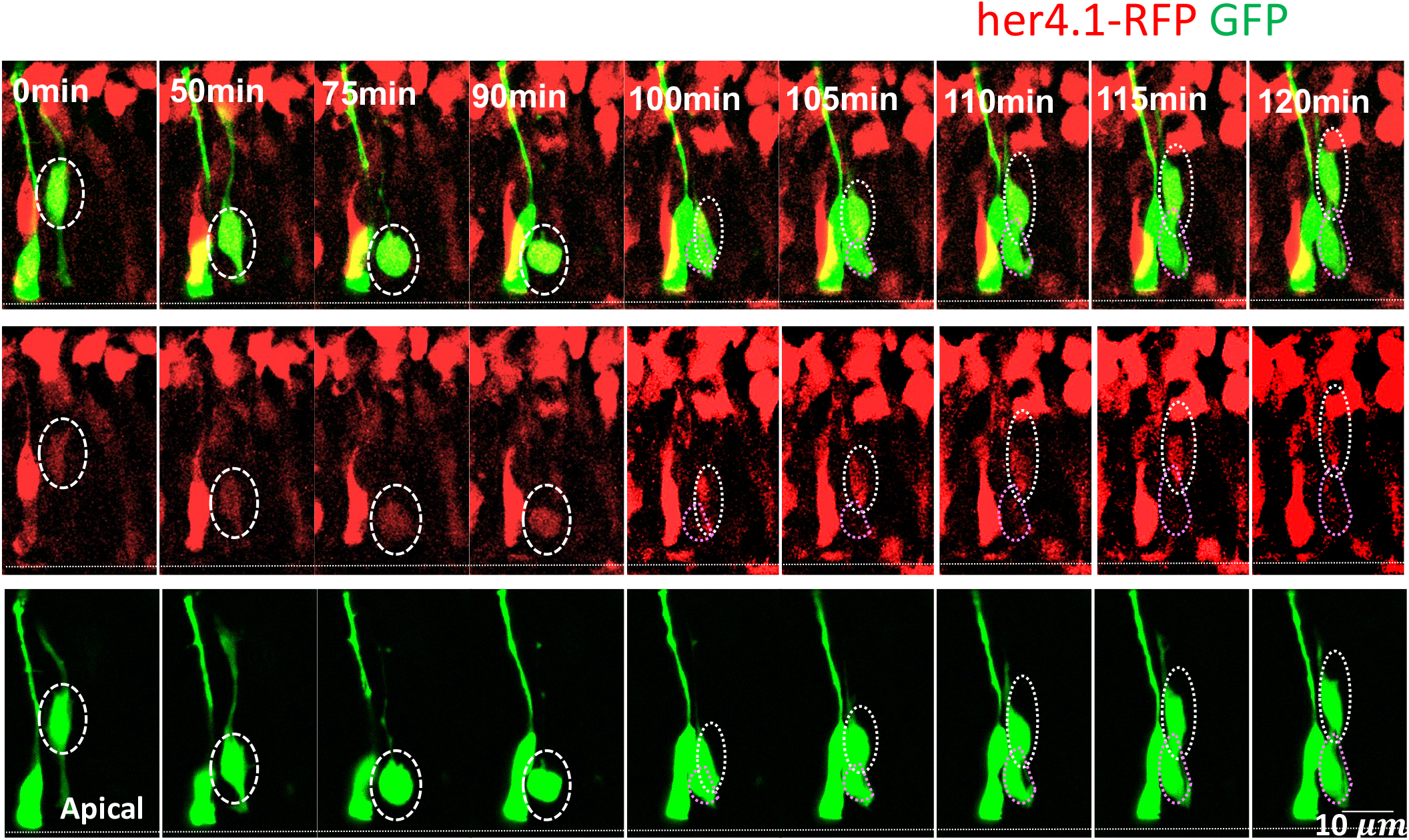
Posterior daughter migrates more basally to become Notch^hi^ in an additional asymmetrically dividing RGP. *In vivo* time-lapse sequence of images showing a sparsely labeled mitotic RGP in *Tg[her4.1-dRFP]* embryo, which divided along the A-P axis; the posterior daughter migrated basally and expressed higher level of her4.1-dRFP (i.e. Notch^hi^).

**Extended Data Figure 6.**
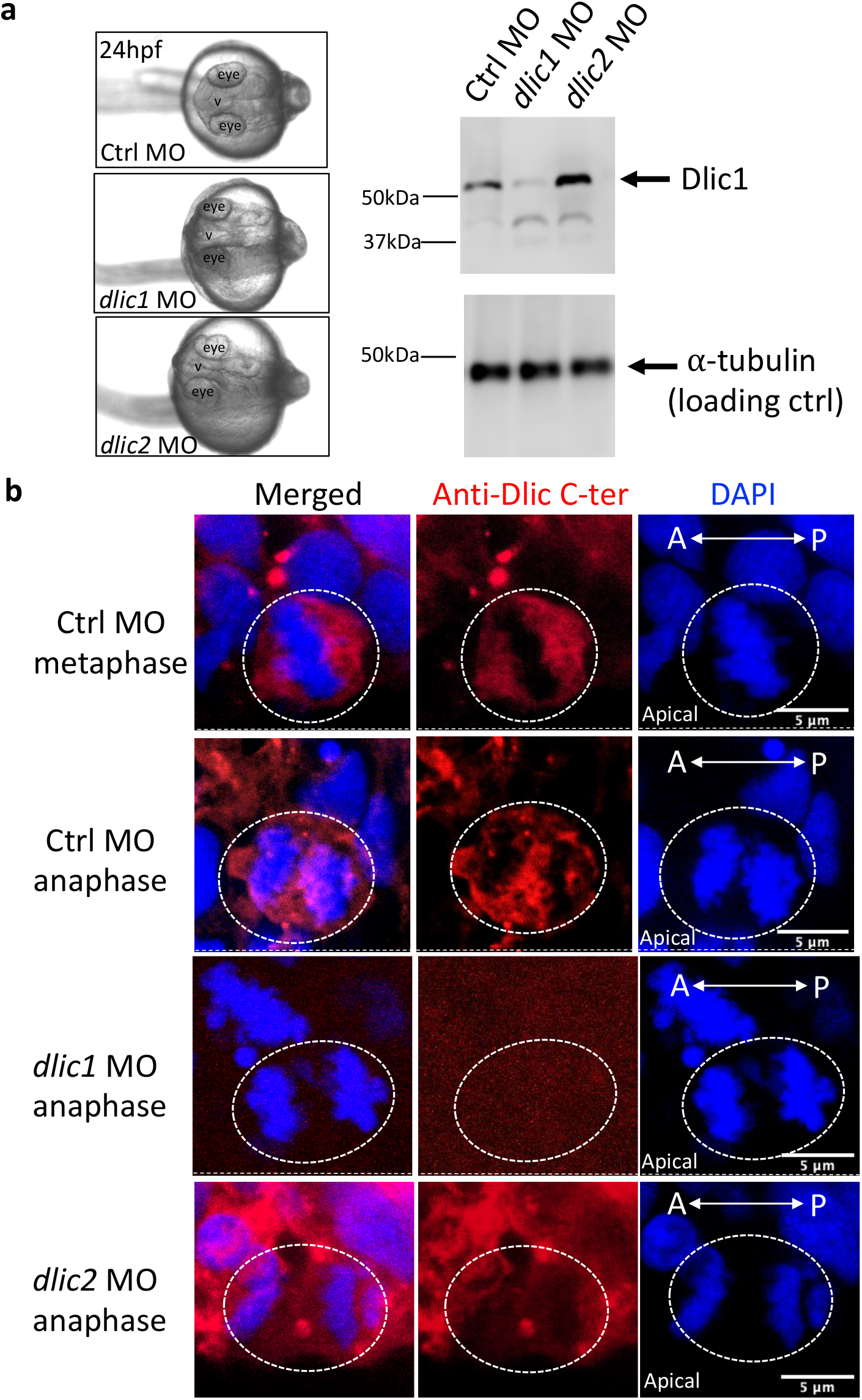
The Dlic1 isoform is predominantly expressed in mitotic RGPs. **a**, Morphological images (left) showing that both *dlic1* and *dlic2* MO knockdown reduced the overall size of embryos at 24 hpf (n=30 embryos analyzed). Western blotting (right) using an anti-*Drosophila* Dlic antibody showing a band at the expected size of ~50 KD. The intensity of the band is significantly decreased in *dlic1* MO but increased in *dlic2* MO, indicating that the antibody recognizes the Dlic1 protein in zebrafish. A small minor band of unknown identity was also detected at ~40 KD. **b**, IF labeling of mitotic RGPs in 28 hpf embryo showing staining with anti-Drosophila DLIC C-ter antibody. Immunoreactivity is observed in control (15 metaphase and 8 anaphase RGPs from 6 embryos of two repeat experiments and *dlic2* MO (6 anaphase RGPs from 4 embryos) but lost in *dlic1* MO (10 metaphase and 5 anaphase RGPs from 5 embryos of three repeat experiments), indicating that the antibody specifically recognizes Dlic1 isoform in mitotic RGPs. Max projections of 20 confocal z-stacks (z-step is 0.26 μm) were shown for each cell.

**Extended data Figure 7.**
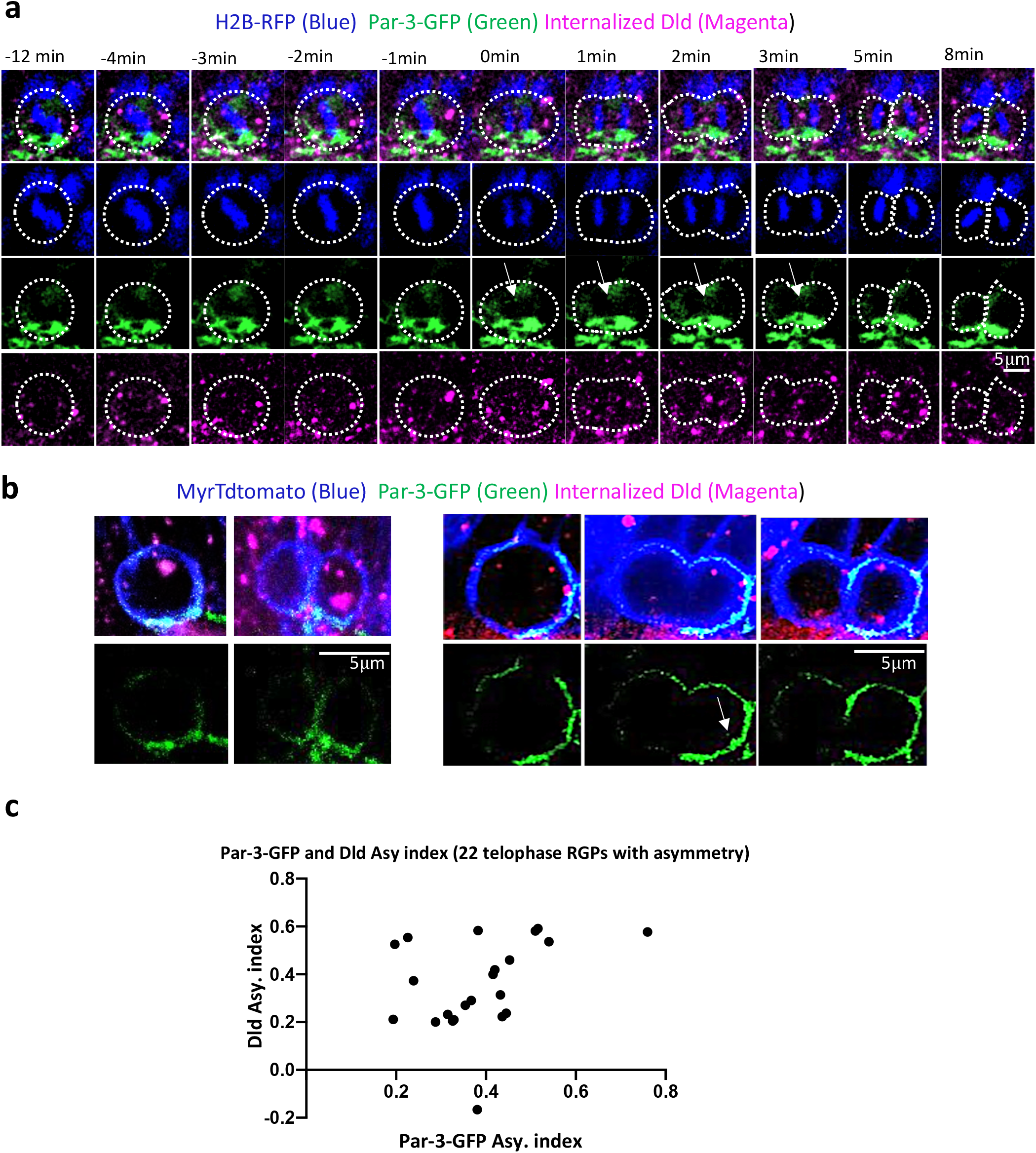
Both Par-3 and Dld endosomes preferentially segregate into the posterior daughter of asymmetrically dividing RGPs. **a**, *In vivo* time-lapse sequence of images showing the dynamics of Par-3-GFP and internalized Dld in asymmetrically dividing RGP. Cytosolic Par-3-GFP was observed, albeit at reduced intensity compared to the membrane form. DNA is labeled with H2B-RFP (pseudo-colored blue). **b**, Montage of images showing two mitotic RGPs with membrane (Myr-TdTomato, pseudo-colored blue) instead of nuclear labeling, together with Par-3-GFP and internalized Dld. **c**, quantification plot showing the correlation of Par-3-GFP and Dld endosome asymmetry indices (See Methods). 22 asymmetrically dividing RGPs at telophase (with enriched posterior Par-3-GFP) were included in the quantification. The correlation coefficient r = 0.34.

**Extended data Figure 8.**
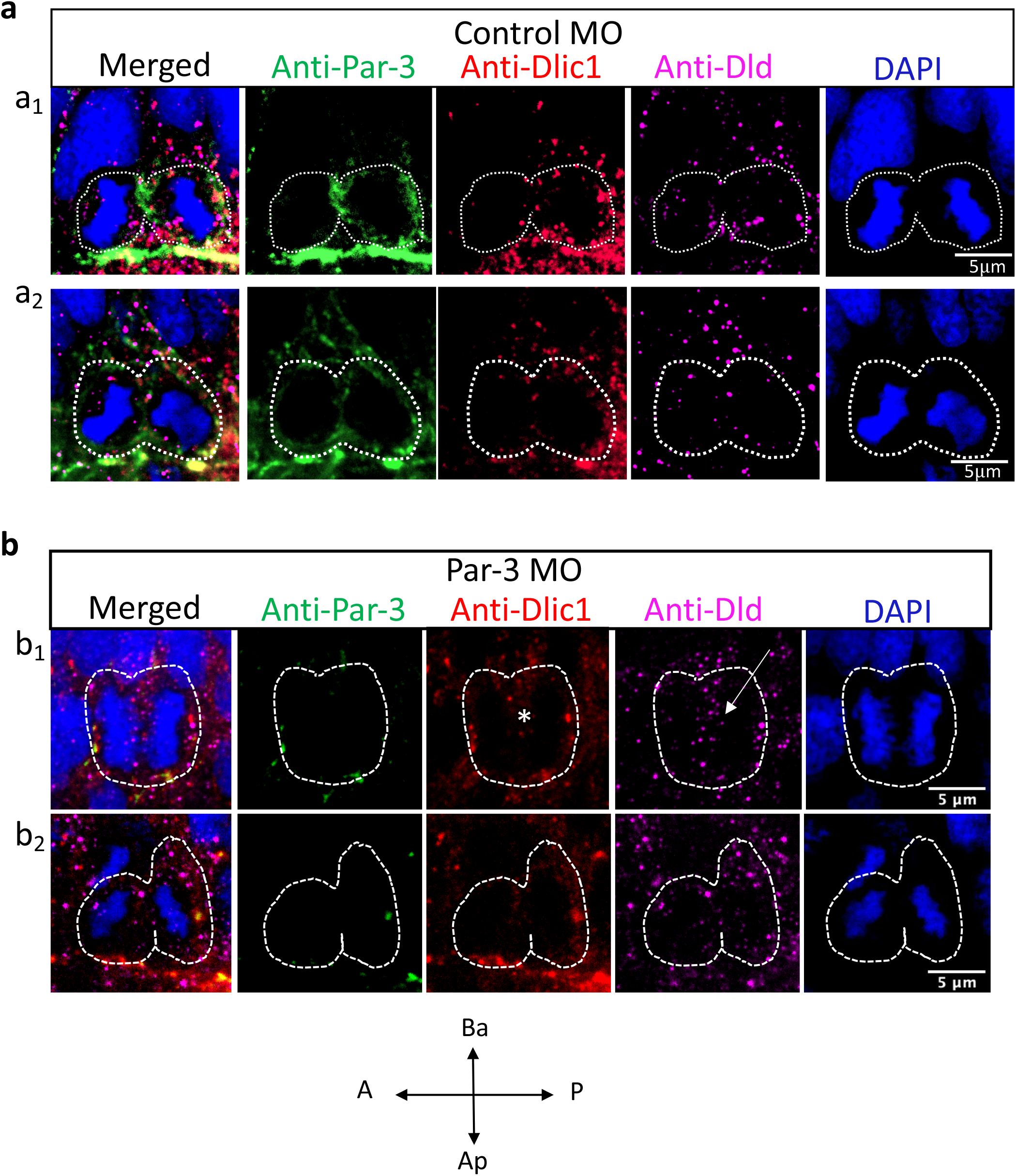
Validation of an anti-Par-3 antibody by immunofluorescence labeling (IF) in mitotic RGPs. **a**, IF labeling with anti-Par-3, anti-Dlic1, and anti-Dld in mitotic RGPs from control MO-injected 28 hpf embryo. Two representative RGPs are shown (a1 and a2). **b**, the same staining in *par-3* MO injected embryos showing the loss of Par-3 immunoreactivity. Similar results were observed in 13 mitotic control RGPs from 6 embryos of 3repeat experiments, and 13 par-3 MO RGPs from 6 embryos of 3 repeat experiments.

**Extended data Figure 9.**
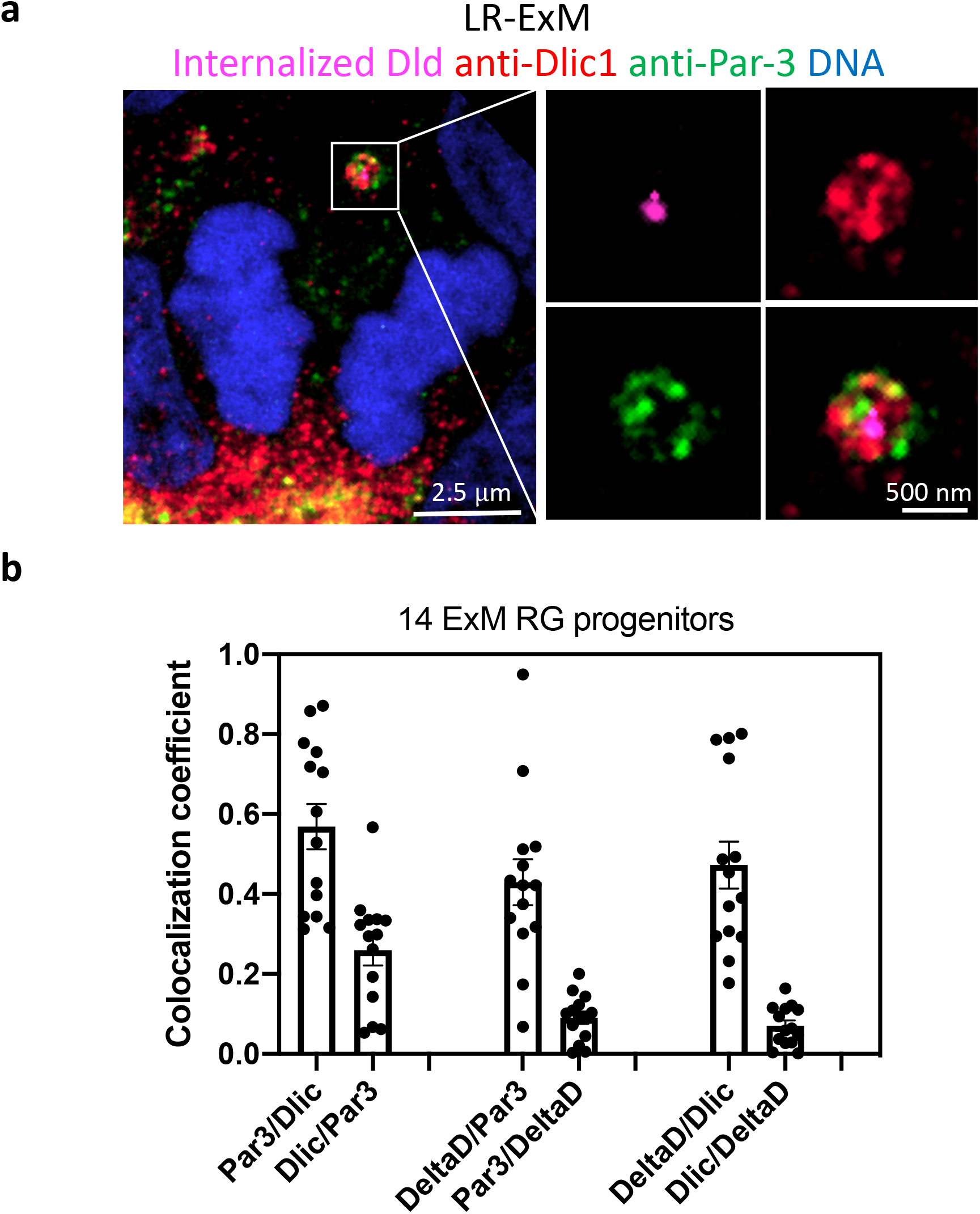
Endogenous Par-3 and Dlic1 decorate Dld endosomes in mitotic RGPs. **a**, Maximum projection of 5 confocal z-stacks showing the co-localization of internalized Dld, anti-Dlic1, and anti-Par-3 on a ring-shaped endosome in anaphase RGP (expanded 4-fold). **b**, Plot showing co-localization co-efficient of anti-Par-3, anti-Dlic1 and internalized Dld fluorescence in 14 RGPs processed by LR-ExM. The results were obtained from 6 embryos of 4 repeat experiments. Mean with S.E.M was shown for each group.

